# Immediate predation risk alters the relationship between potential and realised selection on male traits in the Trinidad guppy *Poecilia reticulata*

**DOI:** 10.1101/2022.04.03.486867

**Authors:** Alexandra Glavaschi, Silvia Cattelan, Alessandro Devigili, Andrea Pilastro

## Abstract

Predation risk perception can alter mating behaviours in males and females, but the consequences for sexual selection remain underexplored. We have previously shown that in experimental populations of Trinidadian guppies *Poecilia reticulata* the opportunity for sexual selection (i.e. the variance in male reproductive fitness) was higher following exposure to a simulated risk of predation than in a no-risk condition. We build upon this result by exploring whether imminent predation risk affects: 1) the relationship between the opportunity for sexual selection and the actual strength of selection on male traits and 2) the traits contributing to male fitness, and he shape of selection on these traits. While predation risk increased the variance in male fitness, realised selection on traits remained unaffected. Pre- and postcopulatory traits follow complex patterns of nonlinear and correlational selection in both treatments. Differences in selection gradients deviate from predictions based on evolutionary responses to predation, the most notable being stronger selection on courtship rate under predation risk. Our results demonstrate that the operation of sexual selection can be altered by perception of an imminent predation risk and reinforce the notion that both trait-based and variance-based metrics should be employed for an informative quantification.

## Introduction

Non-lethal effects of predation are increasingly more often recognised as relevant drivers of changes in prey populations (1–3). Perception of predator cues can affect provisioning of offspring, ultimately determining their fitness (4–6). Other non-lethal effects consist of changes in mating behaviour in both males and females, with the potential to impact sexual selection trajectories. Changes in female choosiness and/or preference for conspicuous male traits following predator threat perception have been reported in variable field crickets *Gryllus lineaticeps* (7), lesser waxmoths *Achroia grisella* (8), and swordtails *Xiphophorus helleri* (9, 10). Exposure to predator cues is associated with a higher number of failed copulations in *Pardosa milvina* wolf spiders (11) and with a reduction in male courtship in *Schizocosa* wolf spiders (12). Predation threat perception can intensify sexual conflict if females engaged in antipredator behaviours are not able to evade unwanted mating attempts(13). For example, female water striders perceiving a risk of predation are more likely to accept matings they would otherwise avoid, particularly with large males (14, 15). Similarly, females of pygmy squid *Idiosepius paradoxus* remove fewer forcibly-inserted spermatangia in the presence of predator cues (16), with possible effects for sperm competition.

Whether or not non-lethal effects of predation do influence the operation of sexual selection (e.g. its strength, shape, targeted traits) remains underexplored (17). This is perhaps surprising, given the increasing awareness of how ecological factors affect sexual selection dynamics (18–21) and the omnipresence of predation risk in the wild. Moreover, immediate predation risk can vary over short time scales irrespective of the background predation intensity characteristic of the habitat (22, 23), therefore it is plausible that the same population experiences consecutive reproductive episodes under different levels of predation risk, with potentially different outcomes for the shape and strength of sexual selection and the traits involved.

One species in which the effects of predation risk on aspects of reproduction are extensively documented is the guppy *Poecilia reticulata*. On its native island of Trinidad, this small freshwater fish with internal fertilisation inhabits rivers and pools along a predation gradient, with consequences for ecology and life-history traits (24, 25). Males from low-predation localities reach maturation at a larger body size, are more brightly-coloured and perform courtship behaviours (sigmoid displays; SDs hereafter) at relatively high frequencies, while males from high-predation populations are smaller, duller and rely more heavily on forced copulation attempts (gonopodial thrusts; GTs hereafter)(24). In addition, predation regime indirectly affects postcopulatory traits linked to guppy sperm performance (26). Observations in the lab indicate that more colourful males with a higher courtship rate are usually preferred by females and have a higher reproductive success (25, 27). In the presence of predator cues, males reduce the frequency SDs while increasing the rate of GTs (28). This change in male mating tactic is partly mediated by a reduction in female receptiveness (29) and preference for conspicuous male colouration (30). At the same time, if given the chance to observe male behaviour in the presence of predators, females prefer bolder males (that show a higher propensity to take risks, i.e. (31)), who indeed benefit from a higher reproductive success compared to their shier counterparts (32).

We have previously shown that simulation of immediate predation risk increases the strength of sexual selection (expressed as the standardised variance in male reproductive success, *I_RS_*) on guppy males (33). This was mainly driven by a higher variance in mating success, suggesting that, at least in our experimental conditions, predation risk may be associated with stronger sexual selection on male precopulatory traits. The variance in male reproductive success, however, does not necessarily represent the realised selection on traits, but rather an estimation of the upper limit of the strength of sexual selection (34–36), and does not distinguish between contributions from male traits and random variation in reproductive success not attributable to sexual selection (37).

Despite intense debate regarding the use of trait-based statistics (such as selection gradients) or variance-based statistics (for example the opportunity of sexual selection) for quantifying sexual selection (38, 39), comparisons between the two methods have been largely based on simulated datasets (40), while empirical tests have yielded mixed results (41–43).

Here we build on our previous findings by aiming to quantify the effects of immediate predation risk on: 1) the relationship between the total opportunity for sexual selection (standardised variance in reproductive success, *I_RS_*) and the actual strength of selection on male traits and 2) the targets and shape of selection. We conduct multivariate selection analyses followed by canonical rotations, focusing on male traits known to contribute to pre and postmating success in the guppy (20, 31, 44–48).

The effect of immediate predation risk on the strength and shape of selection on male traits will be influenced by female choosiness and polyandry, although the exact pattern is not easily predictable. If predation risk causes a decrease in female choosiness, then mating should be more random with respect to male precopulatory traits (such as body size, colouration and courtship behaviour), leading to weaker sexual selection on these traits (49, 50). On the other hand, if polyandry decreases in response to predation risk (as observed in our guppy population), then the potential for selection on male precopulatory traits should increase (51) while the importance of postcopulatory traits for male reproductive fitness should decrease.

Based on observations of guppies in the wild and in the laboratory (see above), and considering the complex patterns of linear and nonlinear selection identified in our population (20, 52), we can predict that the combinations of traits advantaged under perceived predation risk include boldness and GTs, while in control conditions they comprise orange colouration, SDs, gonopodium length, iridescence and GTs. In addition, in accordance with the reduction in female polyandry observed previously (33), we expect that postcopulatory traits would be less relevant for male fitness in the presence of predation compared to control conditions.

## Materials and methods

### (a) Experiment overview

As described in (33), mating trials in the presence and absence of predation risk were carried out in populations consisting of six males and six virgin females (hereafter “replicates”, see below). Males were selected from stock tanks ensuring that, within the same experimental population, they could be individually recognisable by the human observer from colour patterns. The sequence of data collection is presented in supplementary Figure S1. Briefly, males were isolated individually for three days, then subject to two boldness tests (see below), photographed and stripped of sperm to standardize their initial sperm reserves. Five days after photography, males were subject to mating trials in the first treatment, with the second treatment following six days after. We tested a total of 20 male replicates both in the presence and absence of predation cues (i.e. a repeated-measure design), while the groups of females differed between treatments. Fin clips for the purpose of DNA extraction were obtained from males at the end of behavioural observations and from females after they produced a brood. Offspring were euthanised at 24-48h of age and preserved in pure ethanol at −20°C until processing. Following data collection, all adults were released into post-experimental tanks and not reused in further experiments. Predation risk simulation, observations of mating behaviour and paternity assignment are the same as in (33) and also described in the supplementary material.

### (b) Boldness test

We measured boldness using a modified version of the open-field test. Our setup consisted of a white circular arena, 40 cm in diameter filled with water to a depth of 2.5 cm. The arena contained a 3.5 cm diameter refuge in the centre, manufactured from a plastic bottle cap. A 15-W neon light on each side provided illumination. The plain white background and shallow water very likely generates fear in guppies, which is central to boldness measurement (53). The fish was released close to the refuge and its behaviour recorded for 10 minutes with a Panasonic HCV180 video camera mounted 1m above the arena. Two boldness tests, separated by 48h, were performed for each male. The latency to leave the refuge and the total time spent underneath the refuge were scored from videos using BORIS 7.1.3 ((54), http://www.boris.unito.it/pages/download.html). Both behaviours were repeatable (r ≥ 0.3 according to the formula proposed by (55)). For each behaviour, we calculated the average between the two observations and reduced them to a single variable using a principal component analysis. The loading factor of each original variable was 0.94 and the resulting principal component, hereafter referred to as boldness, explained 89% of the total variance.

### (c) Morphology and sperm assays

Males were anesthetised in a bath of MS-222, placed on a grid-lined slide under a dissection microscope equipped with a Canon 450D camera and their left sides photographed. The ejaculates were stripped into a drop of 0.9% saline solution by swinging the gonopodia (intromittent organs) back and forth and applying gentle pressure to the abdomen. In this species, sperm is organised in discrete bundles (spermatozeugmata), each containing ~22000 sperm cells (56). All bundles were photographed for the purpose of sperm counting. Male body area, gonopodium length, area of colouration (orange and iridescent) and sperm number where scored from pictures using ImageJ software (https://imagej.nih.gov/ij/download.html).

Three sperm bundles were placed on a multi-well slide coated with 1% polyvinyl alcohol to prevent sperm cells from sticking to the glass (47) and activated by 3 μl of water containing 150 mM KCl and 2 mg/L bovine serum albumin (57). Sperm velocity was measured using a CEROS sperm tracker (Hamilton-Thorne Research, Beverly, MA, USA) as cells were swimming away from the dissolving bundle. The sperm tracker provides a series of sperm velocity parameters of which we retained VAP (average path velocity) for further analyses (26). Sperm velocity for each male was measured from 295 ± 14.2 (mean ± S.E.) cells.

Sperm viability was measured with a VitalTest kit (Halotech, Spain). Forty sperm bundles were placed in a 0.5 ml Eppendorf tube containing 40 μl saline solution and broken by vortexing for 90 seconds (58). We transferred 6 μl of the resulting mixture into a 0.5 ml Eppendorf tube to which we added 0.5 μl acridine orange, which stains live cells in green, and 0.5 μl propidium iodide which stains dead cells in red. Fluorescent images of the sample were taken with a Leica 5000 B microscope (Leica Microsystems, Wetzlar, Germany) equipped with a digital camera (DFC480; Leica Microsystems, UK). Sperm cells were counted using ImageJ software and viability was calculated as the proportion of live sperm out of the total, from at least 200 cells.

A summary of the phenotypic characteristics of the males used in this experiment is given in the supplementary material (Table S1).

### (d) Statistical analyses

We estimated the relationships between male relative fitness and phenotype using separate multivariate selection analyses (59) for each predation treatment followed by canonical rotations (60). We calculated fitness as the proportion of offspring sired by each male out of the total number of offspring produced within each replicate. We included (*i*) body area, (*ii*) gonopodium length, (*iii*) area of orange colouration, (*iv*) area of iridescent colouration, (*v*) sperm number, (*vi*) sperm velocity, (*vii*) sperm viability, (*viii*) number of sigmoid displays, (*ix*) number of gonopodial thrusts and (*x*) boldness as predictor variables in both models. The sets of males were repeated across the two conditions, therefore the values for boldness, morphological and ejaculate traits are the same for the control and predation treatments, while sexual behaviour was measured during mating trials, therefore values for SDs and GTs differed between treatments. While predation risk was associated with a reduced average courtship rate (33), between-individual differences remained constant (see supplementary material). We standardised response variables to a mean of one and trait values to a mean of zero and standard deviation of one (59).

First, we conducted linear regressions including all trait estimates to obtain linear selection gradients (β). We then fitted second-order regressions including all linear, quadratic and correlational terms to estimate the matrices of nonlinear selection gradients (hereafter referred to as gamma matrices). Statistical packages underestimate quadratic coefficients by 0.5, therefore we doubled these estimates to obtain the correct values (61).

We compared the linear, quadratic and correlational coefficients between treatments with a Monte-Carlo simulation with 10000 iterations. We compared the observed differences (predation – control) in the coefficients with a random distribution of differences obtained by shuffling each male’s reproductive success across treatments. Significance was calculated as the proportion of iterations in which the observed difference exceeded the 95% distribution in the random differences. We used a similar procedure to estimate differences in standardised variance in reproductive success (*I_RS_*; see also (33)), proportion of variance explained by male traits (R^2^ from full quadratic regressions) and total amount of variance explained by traits (*I_RS_* * R^2^). We obtained standard errors of these point estimators with a bootstrap procedure based on 10000 samples.

Interpreting the size and significance of individual coefficients can underestimate the strength of nonlinear selection (62). To overcome this problem, we conducted canonical rotations of the gamma matrices by multiplying them with the matrices of standardised traits (60). Canonical rotations produce new axes of nonlinear selection characterised by loadings of the original traits, similarly to loadings of original variables on principal components obtained by PCA, and identify combinations of traits under selection beyond pairwise comparisons (63, 64). The number of canonical axes obtained is equal to the number of traits included in the analysis (eigenvectors M1 – M10 in each treatment; see below). Each eigenvector has an associated eigenvalue (λ), equivalent to the quadratic selection coefficient along the new axis. The strength of selection (curvature) along each eigenvector is given by its eigenvalue and the shape of selection by its sign, with positive eigenvalues indicating disrupting selection and negative eigenvalues stabilising selection. We also rotated original linear selection coefficients (β) onto the new traits in order to obtain estimates of linear selection along the new axes (θ) (65). We used the permutation procedure proposed by (66) to calculate the significance of each eigenvector. Analyses were conducted with R 4.0.3 (67) and PopTools 3.2 (68) in Microsoft Excel. We visualised fitness surfaces using the ‘Tps’ function of the ‘fields’ package in R (69).

## Results

In a previous paper (33) we demonstrated that the standardised variance in male reproductive fitness is significantly higher in the predation treatment (*I_RS_* = 1.073) compared to control (*I_RS_* = 0.633; Figure 1. See also (33)). Of this total variance, the proportion explained by traits, as estimated by the multiple regression analyses, was significantly higher in the control treatment (R^2^ = 0.709) compared to predation (R^2^ = 0.546; delta ± SE = - 0.162 ± 0.096, *p* = 0.026; Figure 1). By multiplying the standardised variance in reproductive fitness observed in the two treatments by the proportion of variance explained by traits, we obtained an estimate of the strength of overall sexual selection on the male traits considered in this study. We found that sexual selection on traits was higher in the predation treatment (*I_RS_* * R^2^ = 0.586) compared to control (*I_RS_* * R^2^ = 0.449), although this difference was not statistically significant (delta ± SE = 0.138 ± 0.103, *p* = 0.127; Figure 1).

**Figure 1.**
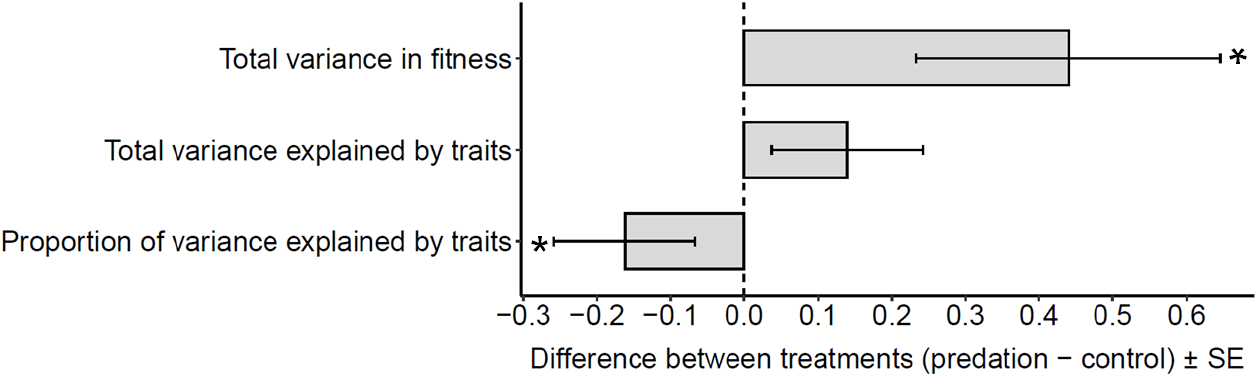
Differences between treatments in standardised variance in fitness (opportunity for selection, *I_RS_*), amount of standardised variance explained by male traits (*I_RS_* * R^2^ from second-order regressions), and proportion of variance (R^2^ from second-order regressions) explained by male traits (white bar). Asterisks indicate significant values (p < 0.05).

When reproductive fitness was analysed separately for each treatment, we did not find any significant linear selection gradients (β) (Table S2). We identified significant quadratic selection on body area (disruptive) and GTs (stabilising) in the control treatment and on SDs (disruptive) in the predation treatment. In the control treatment, all male traits apart from sexual behaviour were involved in significant correlational selection (Table S2), whereas a single negative correlational gradient, between sperm number and boldness, was significant in the predation treatment (Table S2). When we compared the multiple regression coefficients between treatments, we identified significant differences in two linear coefficients (GTs and boldness), two quadratic coefficients (sperm velocity and SDs) and four correlational coefficients associated with sperm number in combination with body area, iridescence, sperm velocity and SDs, respectively. Among these, SDs showed the most pronounced difference between treatments (Table S3). In summary, we identified different predictors of fitness in the two treatments whose coefficients, in turn, are associated with significant between-treatment differences. Thus, in the predation treatment, high and low frequencies of SDs predict high fitness (i.e. disruptive selection), whereas reproductive success in the control treatment is associated with a positive correlation between body area and sperm number and a negative corelation between area of iridescence and sperm number. Canonical rotations produced ten new axes of selection in each treatment (Table 1). A curvature different from 0 (given by the lambda value) indicates significant selection along the respective axis. This was the case for five axes in each treatment (M1, M2, M8, M9, M10). Note that the eigenvectors in each treatment are obtained from separate canonical rotations and are not equivalent, so M1 from the control treatment represents a different axis in canonical space than M1 from the predation treatment. For simplicity, we restrict our discussion to the strongest (highest absolute lambda value) and most significant (lowest *p* value) two axes in each treatment (70). Thus, the highest lambda values in the control treatment corresponded to M1 and M10, which described disruptive and stabilising selection, respectively. Axis M1 was primarily loaded by body area (positive) and sperm number (negative), while axis M10 was mainly loaded by area of iridescence (positive) and sperm viability (negative). The fitness surface defined by these two axes (Figure 3) reveals peaks at extreme values of M1 and average values of M10. The highest peak is associated with small body area and high sperm count in combination with intermediate values for area of iridescence and sperm viability, whereas the lower peak corresponds to males with large body area, low sperm count, and again intermediate values of sperm viability and iridescent colouration (Figure 3). The most significant axes in the predation treatment (with the highest associated lambda values) were M1 and M10, describing disruptive and stabilising selection, respectively (Figure 3). Axis M1 was mainly loaded by SDs (negative) and body area (positive) and axis M10 was primarily described by sperm number (positive) followed by GTs (positive) and boldness (positive). The surface built by these vectors indicates that most successful phenotypes concentrate around extreme negative values of M1 and intermediate values of M10. These males are small, perform SDs at high frequencies and have intermediate values for sperm count, gonopodial thrusts and boldness. A secondary area of high relative fitness, at the positive end of M1 and average values of M10, is associated with intermediate to low sperm count, GT frequency and boldness and also large body area and low SD frequency (Figure 3).

**Figure 2.**
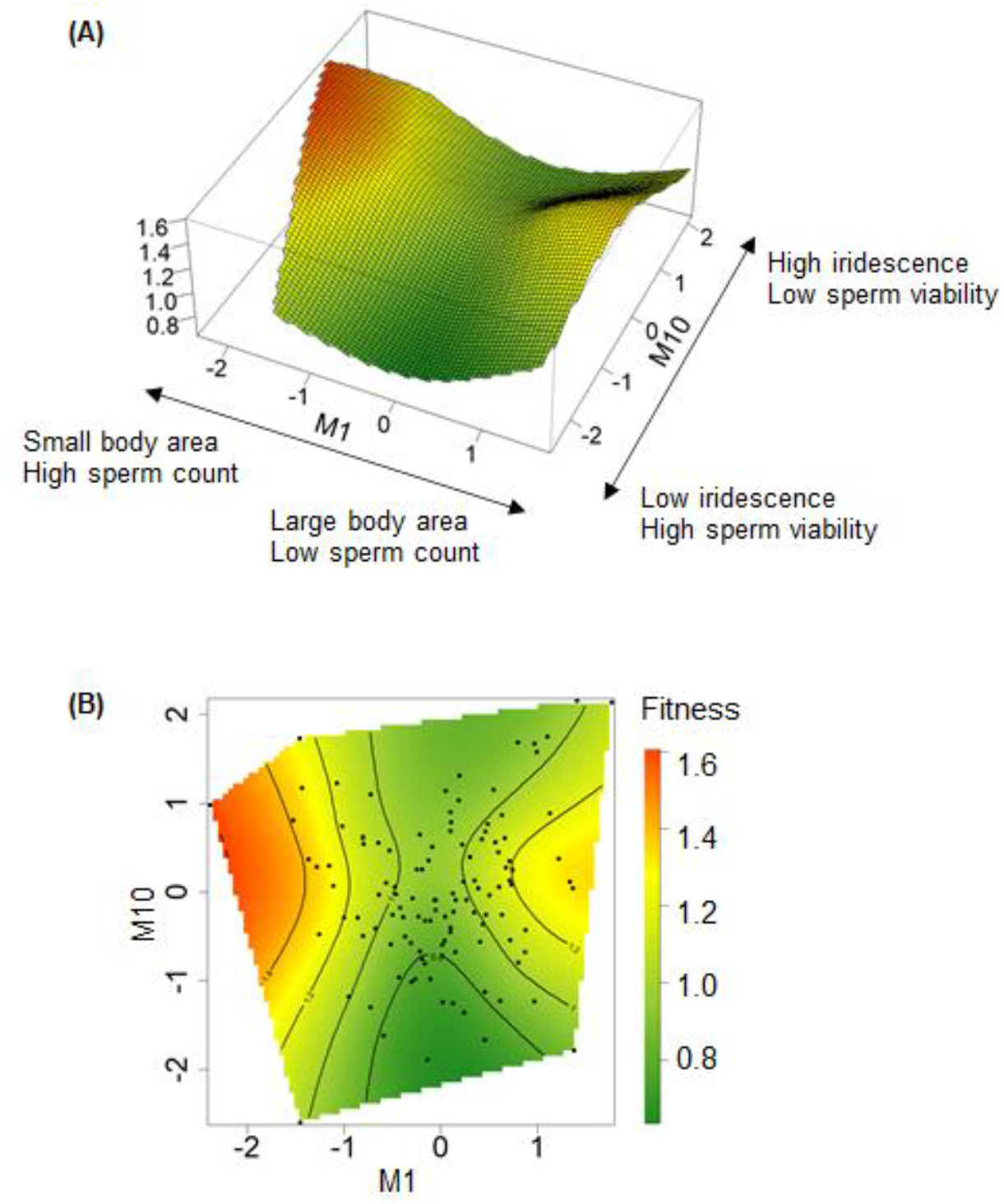
Fitness surface (A) and two-dimensional contour plot (B) illustrating the relationships between relative fitness and major axes of selection in the control treatment. Axis M1 represents disruptive selection and M10 stabilising selection.

**Figure 3.**
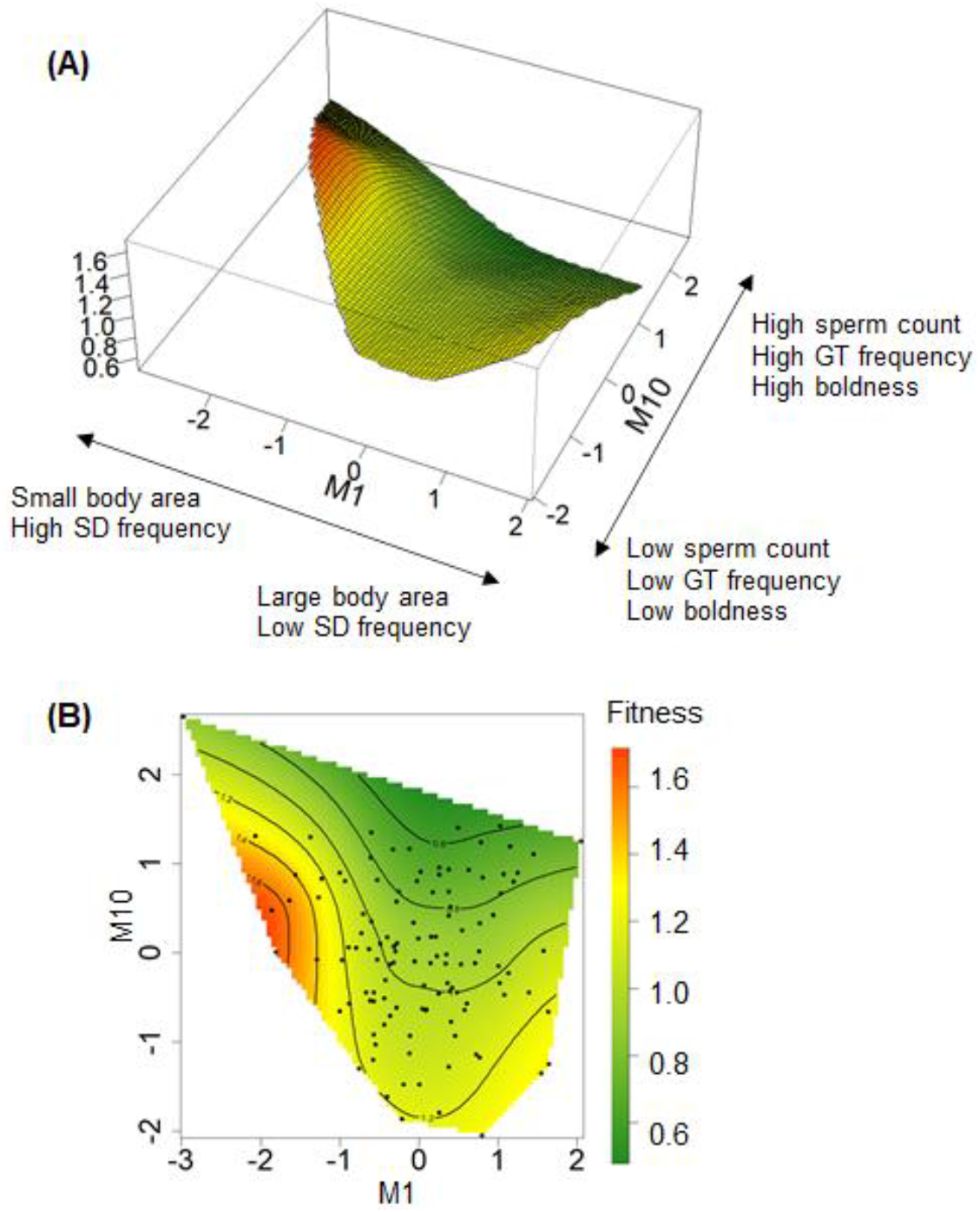
Fitness surface (A) and two-dimensional contour plot (B) illustrating the relationships between relative fitness and major axes of selection in the predation treatment. Axis M1 represents disruptive selection and M10 stabilising selection.

**Table 1.** Eigenvectors obtained by canonical rotations of the gamma matrices and estimates of linear (theta) and nonlinear (lambda) selection gradients along each axis (M1-M10) in each predation treatment. Trait loadings on each eigenvector can be interpreted similarly to those obtained by a principal component analysis. The strength of selection (curvature of the surface) is given by eigenvalues and the shape by their signs (positive=disruptive; negative=stabilising). Significant lambda values (*p* < 0.05) are indicated in bold. *P* values obtained with permutation tests (5000 iterations) following (66).

Control
|  | Theta (P value) | Lambda (P value) | Body area | Gonopodium | Orange | Iridescent | Sperm number | Sperm velocity | Sperm viability | SD | GT | Boldness |
| --- | --- | --- | --- | --- | --- | --- | --- | --- | --- | --- | --- | --- |
| **M1** | -0.038 (0.739) | **2.000 (<0.010)** | 0.694 | -0.020 | -0.171 | -0.277 | -0.536 | -0.157 | -0.284 | -0.071 | 0.079 | 0.093 |
| **M2** | -0.068 (0.414) | **0.854 (0.020)** | -0.040 | -0.071 | -0.416 | -0.281 | 0.212 | 0.412 | -0.382 | 0.116 | 0.129 | -0.593 |
| M3 | 0.102 (0.171) | 0.342 (0.086) | 0.578 | -0.230 | 0.377 | 0.140 | 0.504 | -0.174 | 0.009 | 0.186 | -0.208 | -0.298 |
| M4 | -0.046 (0.533) | 0.204 (0.296) | -0.186 | -0.670 | 0.286 | -0.429 | -0.260 | 0.212 | 0.110 | 0.253 | -0.235 | 0.078 |
| M5 | 0.002 (0.992) | -0.077 (0.690) | -0.360 | -0.196 | 0.052 | -0.133 | 0.062 | -0.712 | -0.496 | -0.193 | -0.045 | -0.126 |
| M6 | -0.019 (0.788) | -0.183 (0.216) | 0.089 | -0.286 | -0.291 | 0.083 | 0.108 | 0.160 | 0.102 | -0.738 | -0.474 | 0.030 |
| M7 | 0.046 (0.583) | -0.338 (0.128) | -0.006 | 0.210 | -0.243 | -0.028 | 0.193 | 0.055 | -0.369 | 0.400 | -0.595 | 0.453 |
| **M8** | 0.028 (0.744) | **-0.585 (<0.010)** | 0.040 | -0.428 | -0.640 | 0.190 | 0.098 | -0.323 | 0.329 | 0.314 | 0.193 | 0.110 |
| **M9** | 0.094 (0.353) | **-0.996 (<0.010)** | 0.095 | -0.011 | 0.033 | -0.544 | 0.533 | 0.031 | 0.030 | -0.199 | 0.387 | 0.468 |
| **M10** | 0.069 (0.445) | **-1.096 (<0.010)** | 0.017 | -0.381 | 0.131 | 0.531 | 0.009 | 0.300 | -0.509 | -0.048 | 0.333 | 0.304 |

Predation
|  | Theta (P value) | Lambda (P value) | Body area | Gonopodium | Orange | Iridescent | Sperm number | Sperm velocity | Sperm viability | SD | GT | Boldness |
| --- | --- | --- | --- | --- | --- | --- | --- | --- | --- | --- | --- | --- |
| **M1** | -0.159 (0.246) | **1.501 (0.010)** | 0.405 | -0.124 | 0.070 | -0.100 | 0.222 | -0.386 | 0.040 | -0.774 | 0.076 | -0.026 |
| **M2** | 0.15 (0.253) | **1.412 (0.014)** | 0.405 | -0.254 | -0.519 | -0.248 | 0.041 | -0.226 | -0.184 | 0.307 | -0.339 | 0.380 |
| M3 | 0.065 (0.584) | 0.679 (0.206) | 0.033 | -0.112 | 0.427 | 0.598 | 0.213 | -0.326 | -0.446 | 0.171 | -0.246 | 0.073 |
| M4 | -0.111 (0.309) | 0.220 (0.597) | 0.021 | 0.245 | 0.231 | 0.076 | -0.556 | -0.341 | 0.287 | -0.002 | 0.105 | 0.601 |
| M5 | -0.041 (0.66) | 0.036 (0.910) | -0.457 | -0.396 | -0.167 | 0.005 | -0.286 | 0.164 | -0.505 | -0.380 | 0.114 | 0.291 |
| M6 | -0.009 (0.946) | -0.034 (0.898) | 0.495 | -0.466 | 0.169 | 0.298 | -0.085 | 0.547 | 0.205 | 0.038 | 0.203 | 0.170 |
| M7 | 0.023 (0.819) | -0.247 (0.202) | -0.048 | 0.323 | 0.092 | 0.006 | 0.193 | 0.446 | 0.077 | -0.308 | -0.656 | 0.343 |
| **M8** | 0.048 (0.629) | **-0.524 (0.040)** | -0.238 | -0.534 | 0.503 | -0.476 | 0.009 | -0.116 | 0.250 | 0.114 | -0.296 | -0.031 |
| **M9** | -0.061 (0.631) | **-0.913 (0.032)** | -0.389 | -0.256 | -0.365 | 0.411 | 0.350 | -0.179 | 0.546 | -0.036 | 0.005 | 0.167 |
| **M10** | -0.173 (0.127) | **-1.715 (<0.010)** | -0.074 | 0.132 | 0.196 | -0.283 | 0.588 | 0.077 | -0.139 | 0.149 | 0.486 | 0.479 |

In summary, in the control treatment sexual selection is nonlinear, in accordance with previous results (20, 52): the fitness surface has a nearly symmetrical saddle shape, with the two peaks at extreme values of M1 of comparable height, suggesting that the alternative phenotypes benefit from similar reproductive success (Figure 3). In contrast, under predation risk sexual selection tends to be more sloped, as the surface built by the two main axes of selection identifies a phenotype (negative extreme of the M1 axis) which represents a main fitness peak, higher than the one at the positive end, suggesting that relatively small males performing a high frequency of SDs are better favoured compared to males showing the reverse combination of traits (Figure 3).

## Discussion

In a previous study (33) we demonstrated that the perception of an imminent predation risk increases the opportunity for sexual selection, as estimated from the standardized variance in male reproductive success (35, 38). This result was due to an increased variance in male mating success and a reduced polyandry (as derived from the number of sires per brood), confirming that polyandry is negatively associated with variance in male reproductive success (51). In the present study, we used the data on male reproductive success to test whether sexual selection differed in strength and shape in response to predation risk. Specifically, we aimed to explore: 1) whether the greater opportunity for (sexual) selection in the presence of predation risk resulted in stronger overall selection on male traits and 2) whether the perception of imminent predation risk affected the importance of male traits and combinations of traits for reproductive success.

Our results suggest that, despite the larger *I_RS_* under imminent predation risk, the overall strength of sexual selection on male traits (expressed as the proportion of the variance explained by traits multiplied by the total variance) did not differ significantly between treatments. This was because in the predation treatment, male traits explained a lower proportion of the total variance in reproductive success compared to the control treatment (Figure 1). Therefore, although predation risk nearly doubled the opportunity for sexual selection, there was no corresponding increase in the strength of sexual selection on male traits. A widespread limitation of studies aiming to quantify selection in small experimental populations (as in the current work) consists of the noise generated by random variation in trait values. Here we overcame this issue by using a repeated measures design (71), therefore our quantification of sexual selection indices under different levels of predation is particularly informative. Our results provide reliable experimental evidence for the theoretical notion that the opportunity for sexual selection (and selection in general) does not necessarily equal realised sexual selection (38, 39, 72).

There are multiple non-mutually exclusive explanations for the observed relationships between the sexual selection metrics we computed in the two treatments. Here we discuss three. First, our result may be explained by the reduced female mating rate under predation risk and the consequent reduction in the contribution of traits under postcopulatory selection towards the variance in male reproductive success (33). Second, female mate assessment could be less accurate under predation risk. Guppies are an extreme example of multiple male ornaments under simultaneous selection by female choice (20, 52). Evaluating complex phenotypes requires time and cognitive effort (73, 74) that may be limited under an imminent predation threat. Therefore, assuming a theoretically preferred male phenotype, “errors” in mate choice could occur more frequently in these conditions. Stochasticity in female choice should reduce the variance in male reproductive success (if female mate choice was purely stochastic the variance in male reproductive success should tend to zero), which contrasts with our observation that predation risk was associated with an increased variance in both male mating and reproductive success (33). A higher variance in male mating success, however, may arise due to a higher importance of mate choice copying, which has been documented in female guppies both in the presence and absence of predator cues (30, 75). In this scenario, the initial choice of the first mating female in each replicate may benefit the first male to mate, irrespective of his phenotype, as suggested by previous results from our population (Morbiato, Cattelan, Pilastro, unpublished data). Thus, the variance in male mating and reproductive success under predation risk would increase (as observed) without affecting the overall strength of sexual selection on male traits, if females tend to copy the choice of other females more frequently under predation risk. Third, the higher portion of unexplained variance in the presence of predation could be a by-product of traits we did not quantify becoming more important for male fitness under these circumstances. Since it is difficult to know whether analyses of the type conducted here capture all components of male reproductive phenotype, this explanation cannot be ruled out.

Our second aim was to test whether predation risk influences the traits that contribute towards male fitness and/or the shape of selection on these traits. Previous analyses (33) indicate that, under predation risk, selection on postcopulatory traits such as sperm number, velocity, and viability should be weaker and selection on precopulatory traits should be stronger. We did find significant differences between the selection gradients under the two conditions (Table S3), but our results partly deviated from this prediction. One important consideration is that the values for morphological traits are the same in both treatments, as they are unlikely to vary substantially over the duration of the experiment (Figure S1) and are not influenced by predation risk. In addition, our interpretation of the relationship between boldness and fitness is based on the assumption that our boldness estimate in standard conditions reflects male propensity to take risks in other contexts, including mating trials. In contrast, male sexual behaviour was recorded during mating trials and is significantly affected by predation risk (33). It is therefore not surprising that the largest difference in selection gradients involved sexual behaviour, although not in the expected direction (Table S3).

In agreement with previous work on the same population of guppies in conditions similar to our control treatment (20, 52), we found that sexual selection was largely correlational and non-linear. While we found no significant linear β regression coefficients or **θ** coefficients on the M vectors in either treatment (Table 1 and Table S2), comparisons between treatments revealed that selection on GTs and boldness was more strongly linear in the predation treatment, and opposite in direction, compared with control (Table S3). At the same time, males with high and low values for body area and intermediate values of GTs were advantaged in the control treatment. In addition, eight combinations of traits were under correlational selection. In agreement with our expectation that postcopulatory traits should be more important for male reproductive success in the absence of predation, all ejaculate traits contributed towards male fitness in the control treatment, in combinations with morphological traits or boldness (Table S1). Canonical rotations confirmed these patterns: phenotypes under strongest selection in the control treatment were characterized by intermediate values for area of iridescence and sperm viability (M10) and either large body area and low sperm number, or small body area and large sperm number (M1, Figure 2).

Extreme (high and low) frequencies of SDs were advantaged under predation risk. We also identified negative correlational selection between boldness and sperm number in the same treatment (Table S2), indicating that bolder males with low sperm count or shy males with high sperm reserves had a higher reproductive success. Canonical rotations confirmed disruptive selection on SDs under predation risk: axis M1 was loaded positively by body area and negatively by SDs, with the highest relative fitness concentrated around the negative extreme (Figure 3). The most advantageous phenotype in the presence of predation risk consisted of small body area, high SD frequency and intermediate sperm number, boldness and GT frequency (Figure 3).

Our results regarding the traits under selection only partly reflect the expected patterns. We did not find a relationship between GTs and male fitness in either treatment, despite a significant difference (yet in the unexpected direction) in linear gradients (Table S3). We did not observe any successful coercive mating but note that observations only covered 50% of the duration of the trials (33), thus we cannot exclude that forced copulations occurred. Even so, their contribution to male reproductive success was most likely limited, given the low insemination success of this mating tactic (76, 77) and considering that females were virgin and therefore expected to be sexually receptive (i.e. to mate cooperatively more often). This was necessary in order to avoid the production of offspring from previously stored sperm that would have biased our measures of male reproductive success, but it has to be noted that in the wild, virgin females are a minority and their mating behaviour may not be representative to that of the population at large (27). In addition, the stronger positive correlation between SDs and reproductive success in the same treatment is surprising, considering that on average males reduced their courtship effort in the presence of predator cues (33). This observation, coupled with the lower polyandry, suggests that cooperative female mating rate is a key determinant of the strength of sexual selection on male traits while sexual conflict plays a minimal role, at least in our population under these experimental conditions.

In conclusion, our study demonstrates that, although imminent predation risk was associated with a higher opportunity for sexual selection and a stronger association between male mating and reproductive success (33), sexual selection on male reproductive phenotype is not significantly stronger and largely similar in shape to that observed in control conditions. The most notable difference in the operation of sexual selection regards the increased relevance of courtship rate under predation risk compared to the control treatment. This is particularly instructive, in our opinion, because it highlights a situation in which males respond to the presence of predator cues by reducing, on average, the frequency of the behaviour (33), yet its importance for male reproductive success increases. Our results therefore demonstrate that non-lethal effects of predation can influence sexual selection trajectories, but in ways that can neither be deduced from lethal effects (e.g. selection against more conspicuous male phenotypes), nor predicted by behavioural responses of males and females to the perception of an imminent predation risk. Finally, our results confirm that, on its own, the variance in male reproductive success is not a sufficiently informative predictor of the strength of sexual selection, at least in polyandrous species (38, 39, 72).

## Ethical note

Our data collection protocol was approved by the University of Padova Institutional Ethical Committee (permit no. 256 /2018).

## Data availability

The dataset is available as supplementary material.

## Author contributions

AG, AP and SC conceptualised the study. AG performed the experiment and collected the data. All authors contributed to analyses. AG led the writing, with contributions from all authors who approved the final version of the manuscript.

## Acknowledgments

We are grateful to graduate students Martina Bonaldi and Marta Guerra for their valuable help with male traits quantification and paternity analyses, respectively.

