## Supplementary material for "Immediate predation risk alters the relationship between potential and realised selection on male traits in the Trinidad guppy *Poecilia reticulata*"

Alexandra Glavaschi^1^, Silvia Cattelan^1,2^, Alessandro Devigili^1^, Andrea Pilastro^1^

^1^ Department of Biology, University of Padova, Via Ugo Bassi 58/B, 35131 Padova, Italy

^2^ Leibniz Institute on Aging - Fritz Lipmann Institute, Beutenbergstraße 11

07745, Jena, Germany

**Supplementary methods**

*(a) Fish maintenance*

Guppies used in this experiment are descendants of a population collected from the Lower Tacarigua river (high predation site) in Trinidad in 2002, maintained in a semi-natural self-sustaining population at the Botanical Garden of the University of Padova. While in the laboratory, fish are kept in 150L tanks, at an approximately equal sex ratio and fed dry food (DuplarinS) and *Artemia salina* nauplii twice daily. New-born are collected monthly from stock tanks and separated into single-sex tanks as soon as sexual traits become visible, to ensure virginity of females.

*(b) Predation risk simulation and mating trials*

Predation risk simulation is described in detail in (1). Briefly, in the predation treatment, the afternoon before the first mating trial, males and females housed separately in single-sex tanks were exposed to predation cues consisting of a predator model (fish bait resembling a pike cichlid *Chrenicichla alta*, 8 cm long) and 2 ml alarm substance (guppy skin extract, (2)). The following morning, males were transferred into the corresponding female replicate tank, 4 ml alarm substance were added, and live behavioural observations started immediately (see below). At the end of the 90 min-long observation, males were moved back to their original tank. These procedures were repeated for two consecutive days, for a total of 180 minutes of observation per replicate. The fish were left undisturbed in the control treatment but were equally manipulated.

The following behaviours were recorded for each male in each treatment: the number of sigmoid displays (SD), gonopodial thrusts (GT), and successful matings, recognisable from rapid postcopulatory jerks performed by the male after sperm transfer. Following mating trials, females were individually isolated into 4L tanks equipped with nurseries until they produced a brood or for a maximum of 50 days. Female shoaling cohesion increased under predation risk and males reduced their courtship rate (1) as observed in wild guppies exposed to natural predators (3, 4), confirming that our setup successfully manipulated fish perception of predation risk.

*(f) Paternity analyses*

The paternity dataset was the same as that was used in (1). Briefly, DNA was extracted from adult fin clips using a salting out procedure (5) and from juvenile whole bodies using a Chelex-100 protocol (6). Two microsatellites (TTA4 (acc. no. AF164205) (7) and AGAT11 (acc. no. BV097141) (8)) where amplified on a Thermo Fisher thermal cycler and the products genotyped by BMR Genomics (Padova, Italy; <https://www.bmr-genomics.it>). Alleles were scored in Geneious 8.1.9 (Biomatters Ltd. http://www.geneious.com) and paternity was assigned using Cervus 3.0.7 (http://www.fieldgenetics.com) with a 95% confidence. Our measure of male reproductive fitness is based on paternity assignment of 1420 offspring from 193 females (predation: N= 99; control: N= 94) and 119 males (Glavaschi et al. 2020).

**Supplementary results**

The number of SD and GT performed by males in the two environmental conditions was significantly correlated (SDs: r=0.31, p=0.0007; GTs: r=0.29, p=0.0014) and the variance in the number of SD and GT did not vary significantly in the two conditions (SDs: control = 0.286; predation = 0.305; GTs: control =0.436, predation =0.491), suggesting that changes in sexual behaviour were consistent across individuals and not amplified by predation risk perception.

**Supplementary figures and tables**

**Figure S1.** Experimental timeline for one replicate. Treatment order was balanced across the samples.

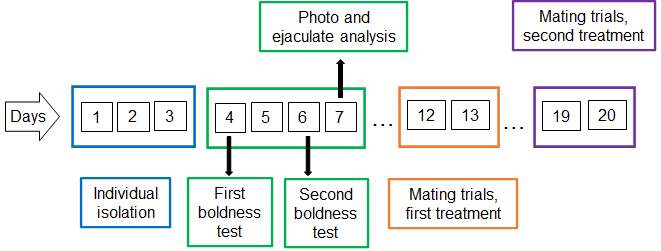

**Table S1.** Average (replicate and total) values ± standard deviations for traits used in selection analyses.

^a^ Boldness is a principal component of proportion of time spent under refuge and latency to leave refuge.

^b^ One male died between treatments therefore this replicate consists of 5 males.

| Replicate | Body area (mm^2^) | Gonopodium  length (mm) | Orange area  (mm^2^) | Iridescent area (mm^2^) | Sperm number  (x 10^6^) | Sperm velocity (VAP; µm/s) | Sperm viability (%) | SD (count) | | GT (count) | | Boldness ^a^ |
| --- | --- | --- | --- | --- | --- | --- | --- | --- | --- | --- | --- | --- |
|  |  |  |  |  |  |  |  | control | predation | control | predation |  |
| MR1 | 71.122  ± 8.369 | 3.731  ± 0.14 | 5.231  ± 2.425 | 10.421  ± 1.957 | 6.098  ± 2.576 | 87.348  ± 2.82 | 0.946  ± 0.047 | 30.667  ± 3.559 | 19.000  ± 13.957 | 4.000  ± 1.414 | 4.167  ± 2.562 | 0.607  ± 0.499 |
| MR2 | 72.413  ± 5.195 | 3.646  ± 0.156 | 6.481  ± 3.871 | 12.535  ± 3.85 | 4.200  ± 4.235 | 95.482  ± 13.323 | 0.981  ± 0.026 | 12.667  ± 9.791 | 29.333  ± 18.457 | 1.833  ± 1.834 | 6.333  ± 1.966 | 0.611  ± 0.425 |
| MR3 | 63.552  ± 6.94 | 3.452  ± 0.217 | 3.962  ± 2.145 | 11.754  ± 2.821 | 3.746  ± 1.827 | 93.069  ± 4.614 | 0.976  ± 0.047 | 13.833  ± 13.014 | 13.500  ± 6.774 | 0.333  ± 0.816 | 3.833  ± 3.371 | 0.938  ± 0.205 |
| MR4 | 70.061  ± 8.568 | 3.469  ± 0.342 | 5.765  ± 1.362 | 13.731  ± 3.028 | 2.407  ± 0.707 | 90.468  ± 7.721 | 0.945  ± 0.064 | 19.000  ± 9.674 | 9.000  ± 7.211 | 6.167  ± 4.75 | 4.500  ± 4.847 | 0.563  ± 0.429 |
| MR5 | 63.613  ± 3.36 | 3.579  ± 0.336 | 3.072  ± 1.954 | 14.382  ± 7.106 | 6.753  ± 3.148 | 88.522  ± 5.871 | 0.905  ± 0.106 | 36.167  ± 18.637 | 14.833  ± 8.084 | 5.500  ± 4.23 | 5.500  ± 2.88 | -0.105  ± 0.993 |
| MR6 | 65.747  ± 9.74 | 3.666  ± 0.172 | 4.991  ± 2.452 | 12.336  ± 3.249 | 7.079  ± 3.528 | 92.017  ± 5.945 | 0.945  ± 0.116 | 27.000  ± 16.149 | 21.500  ± 12.895 | 8.500  ± 6.685 | 6.833  ± 5.307 | -0.156  ± 0.687 |
| MR7 | 68.129  ± 13.352 | 3.475  ± 0.167 | 5.763  ± 3.08 | 12.815  ± 5.04 | 4.492  ± 2.869 | 91.119  ± 5.817 | 0.956  ± 0.057 | 28.333  ± 13.779 | 27.667  ± 15.279 | 7.000  ± 2.529 | 9.833  ± 3.188 | -0.350  ± 1.317 |
| MR8 | 61.015  ± 5.004 | 3.434  ± 0.352 | 5.067  ± 4.128 | 10.943  ± 1.703 | 6.277  ± 2.691 | 104.526  ± 9.804 | 0.984  ± 0.016 | 25.667  ± 16.753 | 24.833  ± 5.98 | 9.167  ± 2.714 | 4.667  ± 2.16 | 0.135  ± 0.9 |
| MR9 | 71.295  ± 10.721 | 3.147  ± 0.484 | 4.785  ± 1.501 | 15.164  ± 3.438 | 7.031  ± 3.372 | 87.572  ± 7.169 | 0.984  ± 0.028 | 32.833  ± 19.964 | 30.167  ± 9.537 | 4.333  ± 1.966 | 5.000  ± 3.794 | 0.090  ± 0.849 |
| MR10 | 59.334  ± 4.222 | 3.188  ± 0.344 | 3.637  ± 3.094 | 10.053  ± 2.709 | 4.324  ± 2.428 | 88.928  ± 4.531 | 0.995  ± 0.008 | 29.000  ± 10.139 | 22.000  ± 5.099 | 5.833  ± 3.868 | 8.000  ± 6.928 | 0.256  ± 0.699 |
| MR11 ^b^ | 56.641  ± 3.34 | 3.290  ± 0.147 | 7.721  ± 4.308 | 8.959  ± 0.444 | 3.327  ± 1.402 | 86.993  ± 9.57 | 0.975  ± 0.023 | 25.800  ± 6.14 | 15.600  ± 3.974 | 8.800  ± 1.643 | 3.200  ± 2.387 | 0.098  ± 0.605 |
| MR12 | 60.678  ± 6.613 | 3.481  ± 0.098 | 4.735  ± 1.725 | 7.650  ± 2.269 | 3.695  ± 2.378 | 94.396  ± 7.361 | 0.986  ± 0.011 | 27.667  ± 11.5 | 24.167  ± 22.373 | 8.333  ± 2.581 | 13.333  ± 6.022 | 0.281  ± 0.563 |
| MR13 | 58.917  ± 3.074 | 3.560  ± 0.23 | 4.732  ± 1.55 | 8.352  ± 1.028 | 4.097  ± 1.98 | 83.684  ± 8.636 | 0.978  ± 0.031 | 29.667  ± 11.927 | 28.667  ± 11.944 | 8.500  ± 3.728 | 11.667  ± 4.76 | 0.322  ± 0.302 |
| MR14 | 62.397  ± 8.337 | 3.282  ± 0.183 | 5.402  ± 1.927 | 10.823  ± 2.183 | 3.867  ± 2.317 | 85.000  ± 8.463 | 0.971  ± 0.054 | 28.333  ± 18.118 | 26.000  ± 13.682 | 8.833  ± 5.528 | 5.833  ± 1.169 | -0.114  ± 0.497 |
| MR15 | 59.467  ± 4.4 | 3.651  ± 0.246 | 3.385  ± 1.261 | 11.157  ± 2.138 | 4.299  ± 1.855 | 74.945  ± 8.982 | 0.737  ± 0.187 | 51.500  ± 16.789 | 23.333  ± 5.006 | 16.167  ± 7.678 | 9.833  ± 4.167 | -0.754  ± 0.694 |
| MR16 | 58.380  ± 2.918 | 3.698  ± 0.166 | 4.357  ± 2.011 | 11.653  ± 1.103 | 4.719  ± 2.684 | 83.668  ± 7.568 | 0.917  ± 0.067 | 24.000  ± 10.139 | 25.833  ± 6.882 | 10.167  ± 4.49 | 4.833  ± 2.483 | -1.219  ± 1.916 |
| MR17 | 56.130  ± 4.237 | 3.386  ± 0.209 | 3.332  ± 1.685 | 10.119  ± 1.41 | 4.317  ± 1.875 | 81.062  ± 2.243 | 0.922  ± 0.054 | 37.333  ± 11.5 | 23.500  ± 11.449 | 10.167  ± 2.926 | 10.000  ± 3.521 | -0.481  ± 1.824 |
| MR18 | 51.385  ± 4.352 | 3.478  ± 0.388 | 3.311  ± 1.186 | 9.630  ± 1.157 | 2.971  ± 2.048 | 82.638  ± 8.251 | 0.868  ± 0.068 | 19.667  ± 10.519 | 25.167  ± 6.765 | 7.500  ± 3.937 | 14.167  ± 4.875 | -0.788  ± 1.255 |
| MR19 | 54.517  ± 5.342 | 3.289  ± 0.522 | 3.408  ± 0.855 | 7.676  ± 3.924 | 2.510  ± 0.443 | 81.248  ± 5.438 | 0.883  ± 0.102 | 19.500  ± 7.687 | 25.333  ± 14.023 | 6.833  ± 2.786 | 10.667  ± 5.391 | 0.142  ± 0.857 |
| MR20 | 64.792  ± 12.824 | 3.524  ± 0.106 | 4.872  ± 1.927 | 13.616  ± 4.33 | 2.356  ± 1.82 | 82.371  ± 8.533 | 0.816  ± 0.401 | 39.833  ± 10.722 | 40.833  ± 19.467 | 11.500  ± 3.271 | 17.000  ± 9.143 | -0.062  ± 0.77 |
| Total | 62.528  ± 8.858 | 3.473  ± 0.303 | 4.675  ± 2.475 | 11.207  ± 3.586 | 4.438  ± 2.704 | 87.759  ± 9.4 | 0.933  ± 0.123 | 27.941  ± 14.936 | 23.580  ± 13.03 | 7.462  ± 4.929 | 8.000  ± 5.604 | 0.000  ± 1 |

| **Control** | Beta coefficients | Gamma coefficients | | | | | | | | | |
| --- | --- | --- | --- | --- | --- | --- | --- | --- | --- | --- | --- |
|  |  | Body area | Gonopodium length | Orange area | Iridescent area | Sperm number | Sperm velocity | Sperm viability | Sigmoid displays | Gonopodial thrusts | Boldness |
| Body area | 0.051 | **1.055** |  |  |  |  |  |  |  |  |  |
| Gonopodium length | -0.011 | -0.027 | -0.183 |  |  |  |  |  |  |  |  |
| Orange | 0.048 | -0.143 | -0.138 | -0.024 |  |  |  |  |  |  |  |
| Iridescent | 0.051 | -0.296 | **0.340** | 0.201 | -0.360 |  |  |  |  |  |  |
| Sperm number | 0.128 | **-0.687** | 0.031 | 0.196 | **0.561** | 0.407 |  |  |  |  |  |
| Sperm velocity | -0.036 | -0.295 | 0.004 | **-0.249** | -0.168 | 0.196 | 0.006 |  |  |  |  |
| Sperm viability | -0.01 | **-0.398** | -0.086 | **0.409** | **0.501** | 0.224 | 0.163 | -0.126 |  |  |  |
| Sigmoid displays | 0.021 | -0.055 | -0.066 | 0.145 | -0.103 | 0.191 | 0.145 | -0.015 | -0.207 |  |  |
| Gonopodial thrusts | 0.023 | 0.031 | 0.244 | -0.174 | -0.069 | -0.254 | -0.038 | -0.023 | 0.049 | **-0.395** |  |
| Boldness | 0.09 | 0.032 | 0.171 | 0.164 | 0.135 | **-0.545** | -0.323 | 0.325 | -0.057 | -0.241 | -0.047 |
| **Predation** |  |  |  |  |  |  |  |  |  |  |  |
|  |  | Body area | Gonopodium length | Orange area | Iridescent area | Sperm number | Sperm velocity | Sperm viability | Sigmoid displays | Gonopodial thrusts | Boldness |
| Body area | 0.034 | 0.295 |  |  |  |  |  |  |  |  |  |
| Gonopodium length | -0.057 | -0.341 | -0.129 |  |  |  |  |  |  |  |  |
| Orange | -0.067 | -0.283 | 0.160 | 0.201 |  |  |  |  |  |  |  |
| Iridescent | 0.008 | -0.141 | 0.096 | 0.699 | -0.066 |  |  |  |  |  |  |
| Sperm number | -0.059 | 0.366 | -0.164 | -0.059 | 0.184 | -0.533 |  |  |  |  |  |
| Sperm velocity | 0.034 | -0.445 | 0.038 | -0.056 | 0.068 | -0.188 | 0.288 |  |  |  |  |
| Sperm viability | -0.071 | 0.124 | 0.338 | 0.185 | -0.330 | -0.129 | 0.218 | -0.127 |  |  |  |
| Sigmoid displays | 0.169 | -0.267 | 0.042 | -0.337 | 0.190 | -0.332 | 0.323 | -0.127 | **0.980** |  |  |
| Gonopodial thrusts | -0.208 | -0.139 | -0.006 | 0.119 | 0.166 | -0.500 | 0.098 | 0.331 | -0.417 | -0.342 |  |
| Boldness | -0.100 | 0.314 | -0.210 | -0.333 | 0.070 | **-0.600** | -0.243 | -0.062 | 0.110 | -0.528 | -0.157 |

**Table S2**. Beta and gamma coefficients for each predation treatments obtained from multiple regressions. Quadratic coefficients are shown on diagonals and indicate disruptive (+) or stabilising (–) selection acting on traits. Correlational coefficients are shown below diagonals and represent pairs of positively or negatively correlated traits under selection. Significant coefficients (*p* < 0.05) are reported in bold.

|  | Coefficient | Predation | Control | Delta | *P* |  | Coefficient | Predation | Control | Delta | *P* |
| --- | --- | --- | --- | --- | --- | --- | --- | --- | --- | --- | --- |
| Linear | Body area | 0.034 | 0.051 | -0.017 | 0.469 | Correlational (continued) | Gonopodium*Sperm viability | 0.338 | -0.086 | 0.425 | 0.158 |
|  | Gonopodium | -0.057 | -0.011 | -0.046 | 0.279 |  | Gonopodium*SD | 0.042 | -0.066 | 0.108 | 0.584 |
|  | Orange | -0.067 | 0.048 | -0.115 | 0.185 |  | Gonopodium*GT | -0.006 | 0.244 | -0.250 | 0.484 |
|  | Iridescent | 0.008 | 0.051 | -0.044 | 0.416 |  | Gonopodium*Boldness | -0.210 | 0.171 | -0.381 | 0.053 |
|  | Sperm number | -0.059 | 0.128 | -0.187 | 0.075 |  | Orange*Iridescent | 0.699 | 0.201 | 0.498 | 0.165 |
|  | Sperm velocity | 0.034 | -0.036 | 0.07 | 0.210 |  | Orange*Sperm number | -0.059 | 0.196 | -0.255 | 0.209 |
|  | Sperm viability | -0.071 | -0.010 | -0.061 | 0.343 |  | Orange*Sperm velocity | -0.056 | -0.249 | 0.193 | 0.104 |
|  | SD | 0.169 | 0.021 | 0.148 | 0.114 |  | Orange*Sperm viability | 0.185 | 0.409 | -0.224 | 0.226 |
|  | **GT** | **-0.208** | **0.023** | **-0.232** | **0.012** |  | Orange*SD | -0.337 | 0.145 | -0.481 | 0.101 |
|  | **Boldness** | **-0.100** | **0.090** | **-0.191** | **0.024** |  | Orange*GT | 0.119 | -0.174 | 0.293 | 0.058 |
| Quadratic | Body area | 0.295 | 1.055 | -0.759 | 0.144 |  | **Orange*Boldness** | **-0.333** | **0.164** | **-0.498** | **0.031** |
|  | Gonopodium | -0.129 | -0.183 | 0.054 | 0.444 |  | **Iridescent*Sperm number** | **0.184** | **0.561** | **-0.376** | **0.002** |
|  | Orange | 0.201 | -0.024 | 0.224 | 0.126 |  | Iridescent*Sperm velocity | 0.068 | -0.168 | 0.236 | 0.093 |
|  | Iridescent | -0.066 | -0.360 | 0.294 | 0.151 |  | **Iridescent*Sperm viability** | **-0.330** | **0.501** | **-0.831** | **0.029** |
|  | Sperm number | -0.533 | 0.407 | -0.940 | 0.062 |  | Iridescent*SD | 0.190 | -0.103 | 0.293 | 0.116 |
|  | **Sperm velocity** | **0.288** | **0.006** | **0.282** | **0.018** |  | Iridescent*GT | 0.166 | -0.069 | 0.235 | 0.125 |
|  | Sperm viability | -0.127 | -0.126 | 0.000 | 0.604 |  | Iridescent*Boldness | 0.070 | 0.135 | -0.064 | 0.425 |
|  | **SD** | **0.980** | **-0.207** | **1.187** | **0.012** |  | **Sperm number*Sperm velocity** | **-0.188** | **0.196** | **-0.384** | **0.017** |
|  | GT | -0.342 | -0.395 | 0.053 | 0.686 |  | Sperm number*Sperm viability | -0.129 | 0.224 | -0.353 | 0.150 |
|  | Boldness | -0.157 | -0.047 | -0.110 | 0.357 |  | **Sperm number*SD** | **-0.332** | **0.191** | **-0.523** | **0.003** |
| Correlational | Body area*Gonopodium | -0.341 | -0.027 | -0.314 | 0.250 |  | Sperm number*GT | -0.500 | -0.254 | -0.246 | 0.751 |
|  | Body area*Orange | -0.283 | -0.143 | -0.140 | 0.335 |  | Sperm number*Boldness | -0.600 | -0.545 | -0.054 | 0.580 |
|  | Body area*Iridescent | -0.141 | -0.296 | 0.155 | 0.435 |  | Sperm velocity*Sperm viability | 0.218 | 0.163 | 0.054 | 0.307 |
|  | **Body area*Sperm number** | **0.366** | **-0.687** | **1.053** | **0.001** |  | Sperm velocity*SD | 0.323 | 0.145 | 0.178 | 0.069 |
|  | Body area*Sperm velocity | -0.445 | -0.295 | -0.149 | 0.159 |  | Sperm velocity*GT | 0.098 | -0.038 | 0.136 | 0.356 |
|  | Body area*Sperm viability | 0.124 | -0.398 | 0.522 | 0.228 |  | Sperm velocity*Boldness | -0.243 | -0.323 | 0.080 | 0.578 |
|  | Body area*SD | -0.267 | -0.055 | -0.212 | 0.491 |  | Sperm viability*SD | -0.127 | -0.015 | -0.111 | 0.605 |
|  | Body area*GT | -0.139 | 0.031 | -0.169 | 0.099 |  | Sperm viability*GT | 0.331 | -0.023 | 0.354 | 0.655 |
|  | Body area*Boldness | 0.314 | 0.032 | 0.282 | 0.248 |  | Sperm viability*Boldness | -0.062 | 0.325 | -0.387 | 0.084 |
|  | Gonopodium*Orange | 0.160 | -0.138 | 0.298 | 0.077 |  | SD*GT | -0.417 | 0.049 | -0.466 | 0.081 |
|  | Gonopodium*Iridescent | 0.096 | 0.340 | -0.244 | 0.413 |  | SD*Boldness | 0.110 | -0.057 | 0.168 | 0.587 |
|  | Gonopodium*Sperm number | -0.164 | 0.031 | -0.194 | 0.171 |  | GT*Boldness | -0.528 | -0.241 | -0.287 | 0.579 |
|  | Gonopodium*Sperm velocity | 0.038 | 0.004 | 0.034 | 0.149 |  |  |  |  |  |  |

**Table S3.** Linear and multiple regression gradients comparison between predation treatment. Delta represents the difference between coefficients from the predation treatment and their corresponding control coefficients. *P* values obtained from Monte-Carlo permutations with 10000 iterations and represent the number of times observed delta is higher than permuted delta. Significant differences (*p* < 0.05) are indicated in bold.
